# The Global State of Genome Editing

**DOI:** 10.1101/341198

**Authors:** Geoffrey H. Siwo

## Abstract

Genome editing technologies hold great promise in fundamental biomedical research, development of treatments for animal and plant diseases, and engineering biological organisms for food and industrial applications. Therefore, a global understanding of the growth of the field is needed to identify challenges, opportunities and biases that could shape the impact of the technology. To address this, this work applies automated literature mining of scientific publications on genome editing in the past year to infer research trends in 2 key genome editing technologies-CRISPR/Cas systems and TALENs. The study finds that genome editing research is disproportionately distributed between and within countries, with researchers in the US and China accounting for 50% of authors in the field whereas countries across Africa are underrepresented. Furthermore, genome editing research is also disproportionately being explored on diseases such as cancer, Duchene Muscular Dystrophy, sickle cell disease and malaria. Gender biases are also evident in genome editing research with considerably fewer women as principal investigators. The results of this study suggest that automated mining of scientific literature could help identify biases in genome editing research as a means to mitigate future inequalities and tap the full potential of the technology.

## Introduction

Genome editing is one of the fastest growing areas in biomedical research and has a high potential in revolutionizing fundamental biological research, engineering of organisms for synthetic applications and eventually clinical medicine [1]. Given the potential societal impact of genome editing technologies, the ethical issues it raises and the economic potential through biomedical applications as well as creation of new jobs, a global understanding of the growth of the field is needed to identify challenges, opportunities and biases that could shape the impact of the technology [2, 3].

Several approaches to genome editing exist-meganucleases [4], Zinc Finger Nucleases (ZFNs) [5], TALENs (Transcription Activator-Like Nucleases) and CRISPR/Cas systems (Clustered Regularly Interspaced Short Palindromic Repeats/CRISPR associated protein) [6]. However, TALENs and the RNA-guided programmable endonucleases -CRISPR/Cas systems- have become the most popular due to the relative ease with which they can be programmed to target DNA in a sequence-specific manner. As genome editing is still in its early days, there may be opportunities to direct its development in a way that maximizes the power of the technology equitably and in a socially responsible manner. In this work, we apply automated mining of abstracts of genome editing publications in 2017 and part of 2018, focusing on the 2 main genome editing technologies-CRISPR/Cas and TALENs to infer the global distribution of genome editing research as a first step to identify emerging biases.

## Results

### General publication trends in genome editing

To obtain scientific publications on genome editing by CRISPR/Cas systems and TALENs, an automated search on PubMed followed by a download of abstracts was performed in the R programming language as described in the Methods section. In 2017, there were 1521 papers directly mentioning CRISPR and genome editing in their abstracts compared to 684 papers for 2018 (up to May 21^st^ 2018). These studies were published across 529 and 349 scientific journals in 2017 and 2018, respectively. In contrast for TALENs based genome editing, there were 146 articles in 2017 as compared to 46 articles in 2018 across 103 and 36 journals, respectively.

The top 5 journals for CRISPR genome editing publications were *Scientific Reports* (81 articles, open access), *Methods in Molecular Biology* (49, subscription), *Nature Communications* (32, open access), *Nature* (30, subscription) and *PLoS ONE* (27, open access) in 2017. The top journals in 2018 were *Methods in Molecular Biology* (24, subscription), *Scientific Reports* (15, open access), *ACS Chemical Biol*ogy (13, subscription), *Nature Communications* (13, open access) and *PLoS ONE* (13, open access). For TALENs, the top journals were *Methods in Molecular Biology* (12 articles, subscription), *Frontiers in Plant Science* (6, subscription), *PLoS ONE* (5, open access), *Developmental Biology* (4, subscription), *Molecular Therapy-Methods & Clinical Development* (4, subscription). A complete table of the number of articles per journal is available in the Supplementary Table 1 and 2. There was a positive correlation between the number of articles published on CRISPR and TALENs (Spearman correlation, *r* = 0.53, *P* = 1.5e-08). A small fraction of journals accounted for a large proportion of most genome editing articles (Figure 1). Specifically, the top 50 journals for CRISPR genome editing in 2017 accounted for 48% of the articles (719 out 1521 articles). These journals constitute only 9.4% (50 out of 529 journals) of the total journals that published at least one article on CRISPR genome editing. A similar observation was made for publications on TALENs where 64% of the articles (93 out of 146 articles) were accounted for by 49% of the journals (50 out of 103 journals). These results show that genome editing publications are disproportionately distributed in a few journals including journals that require a subscription.

**Figure 1:**
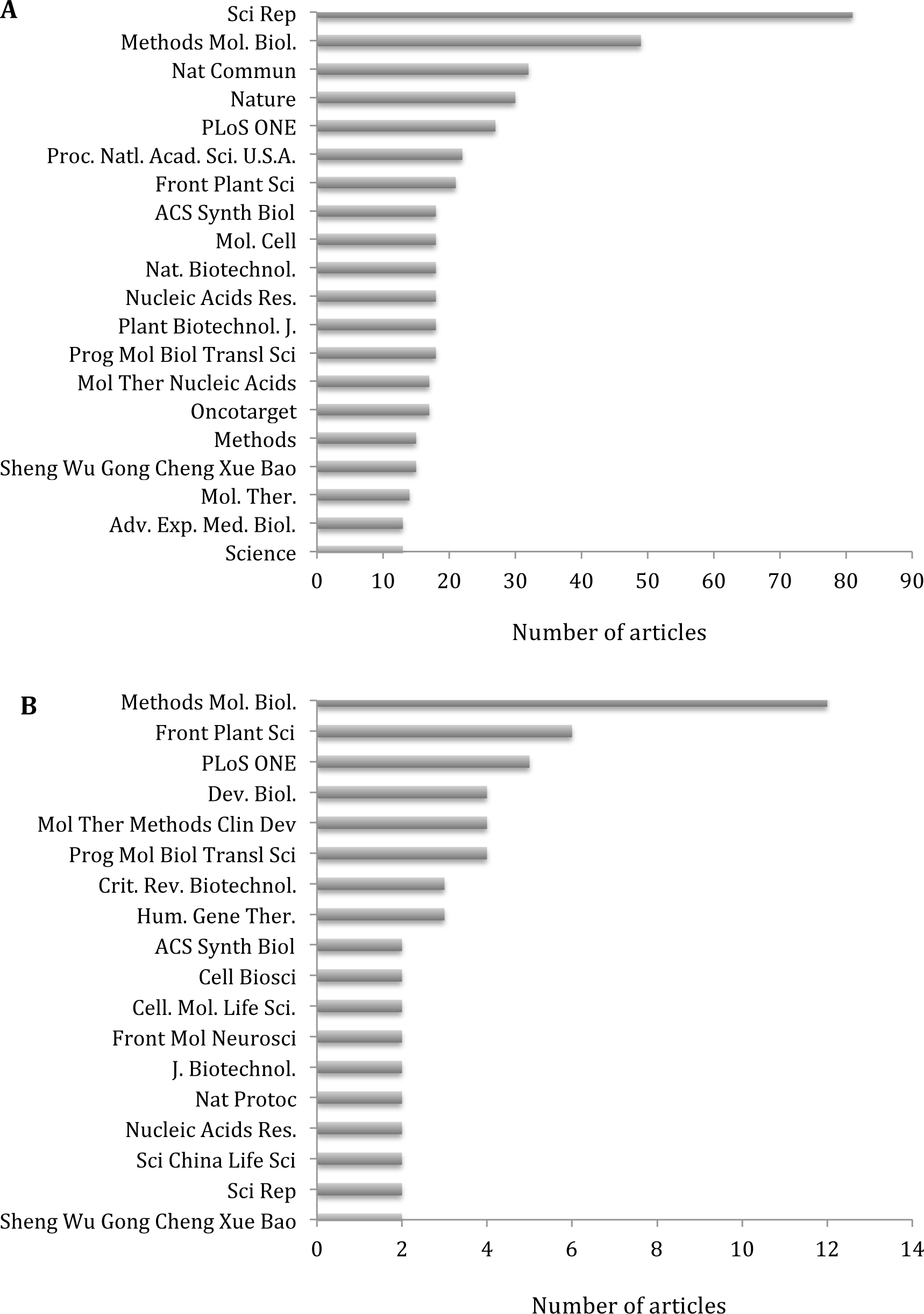
Number of articles on genome editing involving CRISPR/Cas **(A)** or TALENs **(B)** based on the top 50 journals. Journals are ranked by the number of articles mentioning CRISPR genome editing **(A)** or TALENs genome editing **(B)** in their abstracts.

**Figure 2:**
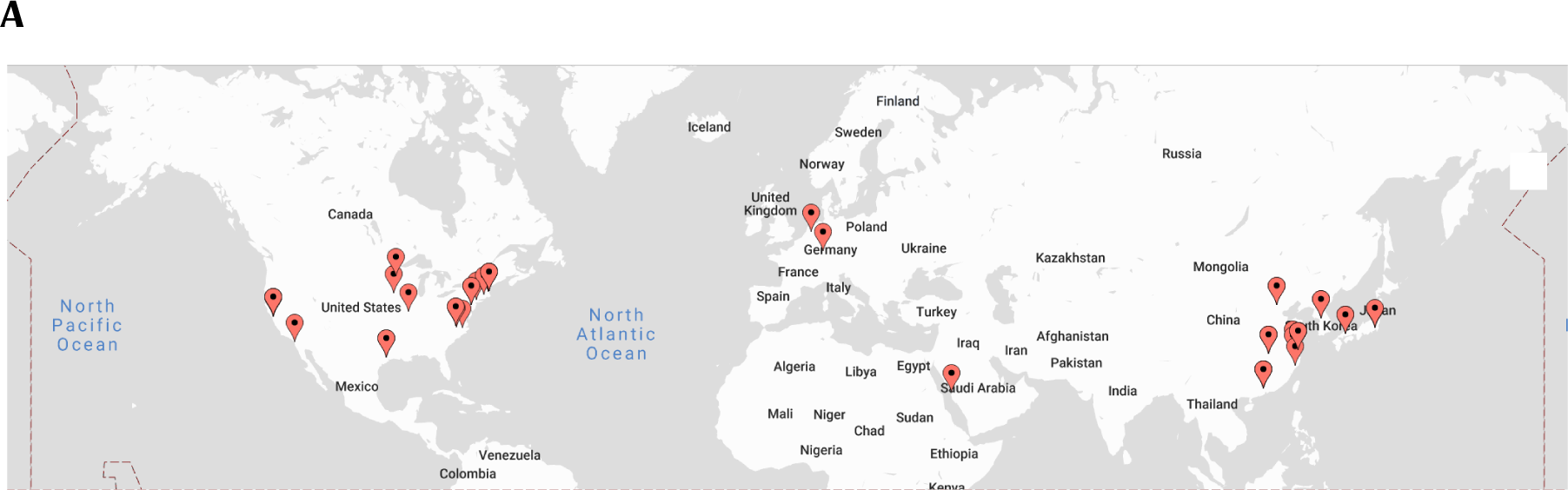
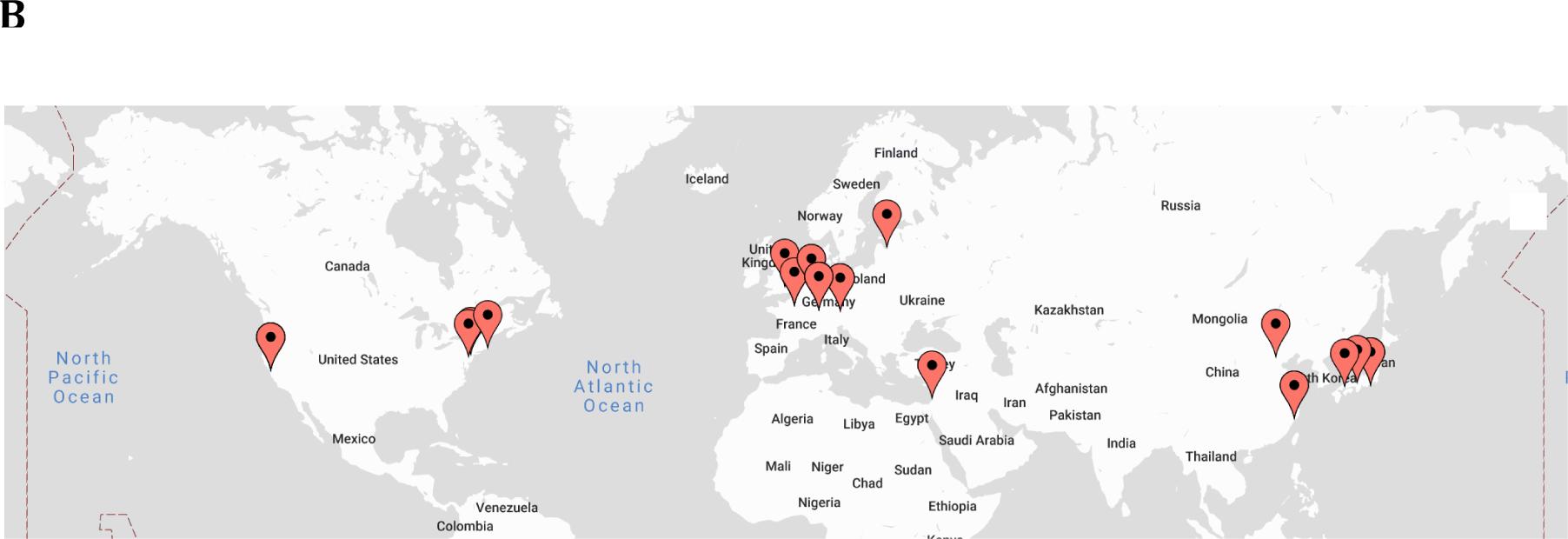
The global distribution of the top 50 authors based on the number of articles published by each author on CRISPR **(A)** or TALENs **(B)** based genome editing in 2017.

### Geographical distribution of genome editing studies

To provide an assessment of the global network of scientists performing genome editing research and publishing their findings, author names were extracted from the downloaded abstracts for the year 2017. In order to identify unique authors, the first names and last names of each author were combined. This resulted in 8,528 unique authors on CRISPR genome editing and 745 authors for TALENs. For CRISPR, the top authors included Feng Zhang (Harvard University, USA; 15 articles), Kim-Jin Soo (Seoul National University, Korea; 15 articles), Gao Caixia (Chinese Academy of Sciences, China; 11 articles), Jennifer Doudna (UC Berkley, USA; 9 articles), Gersbach Charles (Duke University, USA; 9 articles) and Musunuru Kiran (University of Pennsylvannia, USA; 9 articles). Notably, Feng Zhang and Jennifer Doudna who are both central pioneers in CRISPR based genome editing technologies feature in the top authors. In comparison, top authors for TALENs were Sakuma Tetsushi (Hiroshima University, Japan; 5 articles), Yamamoto Takashi (Hiroshima University, Japan; 5 articles), Cathomen Toni (University of Freiberg, Germany; 4 articles) and Mussolino Claudio (University of Freiberg, Germany; 4 articles). It is interesting to note that the top authors for CRISPR genome editing are mainly from the US but they do not feature among the top authors for TALENs. The top 50 authors account for about 20% (301 out of 1521) of CRISPR genome editing publications and 60% of TALENs publications (88 out of 146). The top authors and institutional affiliations are provided in Supplementary Table 3 and 4.

US and China account for approximately 50% of all 8,528 authors for CRISPR genome editing studies in 2017 with 2342 and 2140 authors, respectively. These two countries also lead in terms of the number of authors publishing on TALENs genome editing. Notably, while US leads on CRISPR, China leads on TALENs. The top 20 countries based on the number of authors on CRISPR or TALENs genome editing are provided in Figure 3.

**Figure 3:**
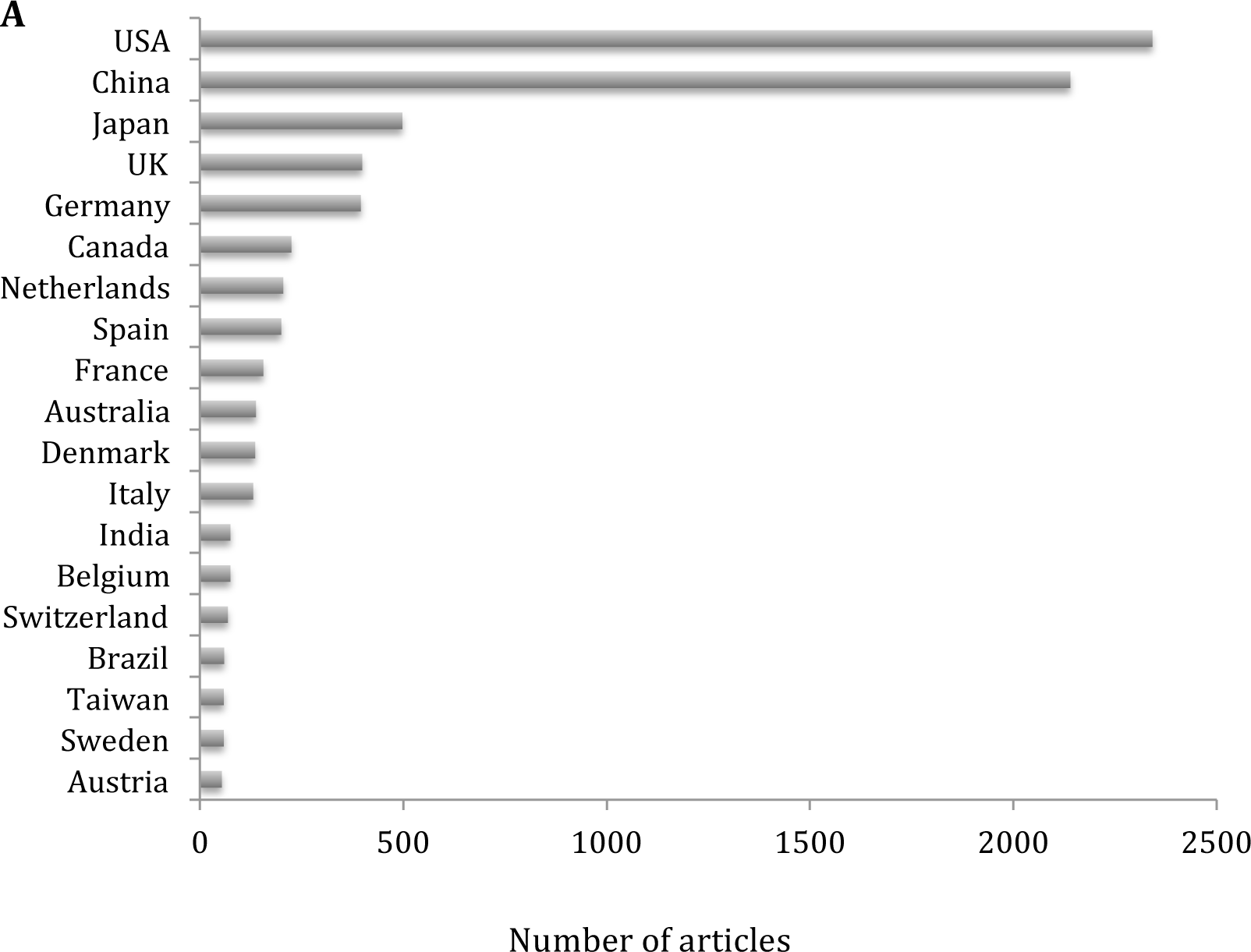
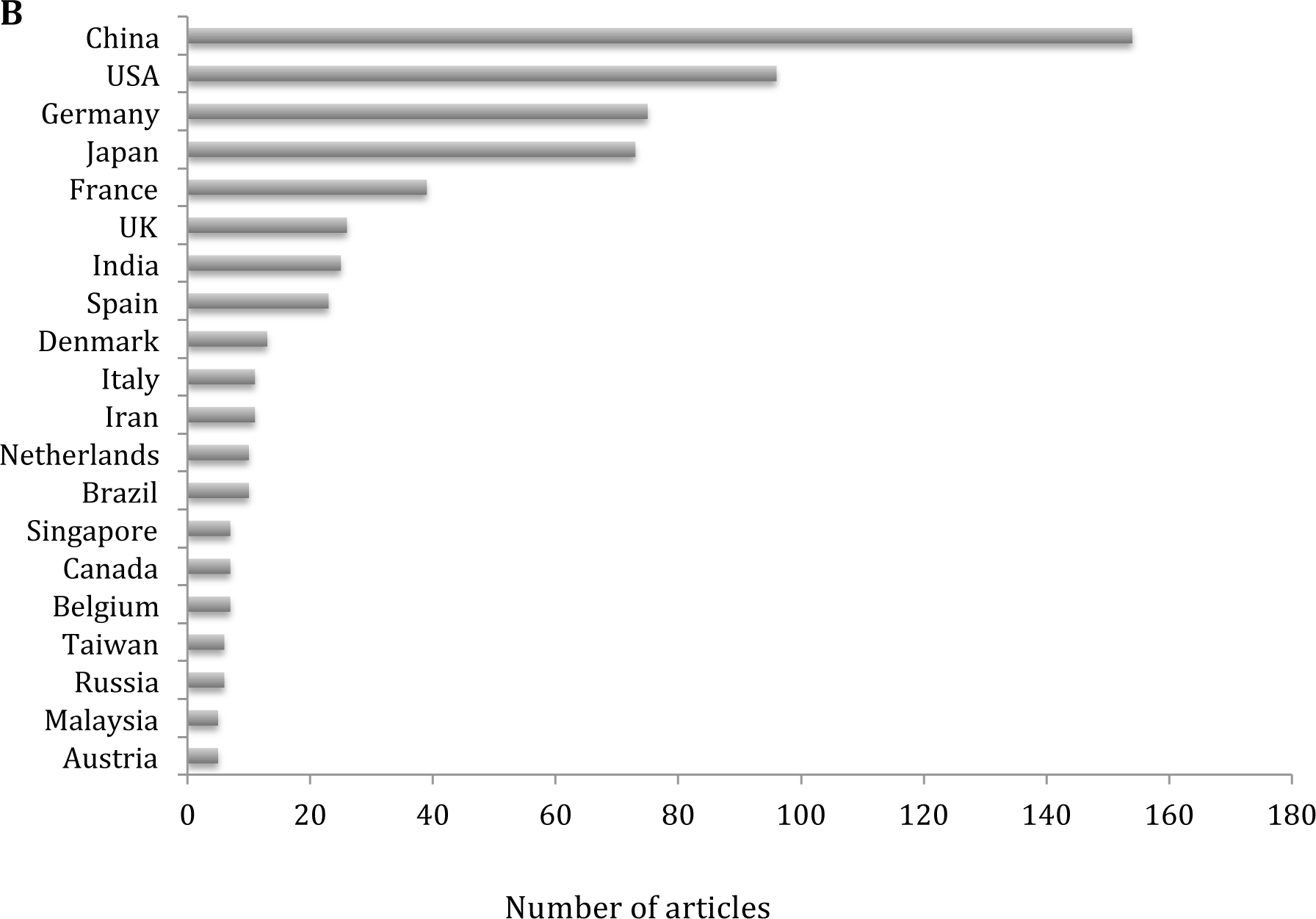
A ranking of the top 20 countries based on number of authors affiliated with institutions in those countries. **(A)** CRISPR based genome editing and **(B)** TALENs based genome editing.

### Gender composition of authors in genome editing studies

Gender diversity is critical in advancing scientific discovery and overcoming health disparities. Yet, women are underrepresented in multiple fields of science and technology [7, 8]. Assessing gender diversity in emerging areas of technology could provide opportunities for addressing disparities before they become entrenched in society. To estimate the gender representation of male and female authors, the first names of authors was used to assign gender based on data from the US Social Security Administration baby names in the R package *‘gender’*. In 2017 for CRISPR genome editing, 43% of first authors were females while 57% were males. In contrast, when examining the last authors, conventionally the principal investigators, only 26% were females and the rest (74%) were males. The disparity between male and female authors was slightly lower when considering all authors of CRISPR genome editing papers irrespective of their positions in the author list (40% were females and 60% males). Authorship of papers on TALENs showed similar patterns: females were 42% of first authors, 28% of last authors and 40% of all authors. These results suggest that pre-existing gender biases in science and technology could be carried over to emerging technologies like genome editing.

### Genes, diseases and species targeted by genome editing studies

Identification of genes and diseases being investigated by genome editing technologies could help in the assessment of potential early clinical applications of the technology and highlight areas that may require new investment. Screening of the downloaded abstracts for gene names led to the retrieval of genes that are the focus of various genome editing studies. Genes were then ranked by the number of genome editing studies in which they appeared in the abstracts. The dystrophin gene (DMD) was the most frequently studied gene using CRISPR, followed by several cancer associated genes including P53 [9, 10], Cystic Fibrosis Transmembrane Conductance (CFTR) gene [11, 12], CXCR4- an HIV-1 entry co-receptor [13, 14]- and the hemoglobin B gene (HBB) important in sickle cell anemia and other hemoglobinopathies (Table 2) [15, 16]. CD4, an HIV-1 receptor [17] was the most studied gene for TALENs genome editing although there were only 2 articles.

**Table 1:**
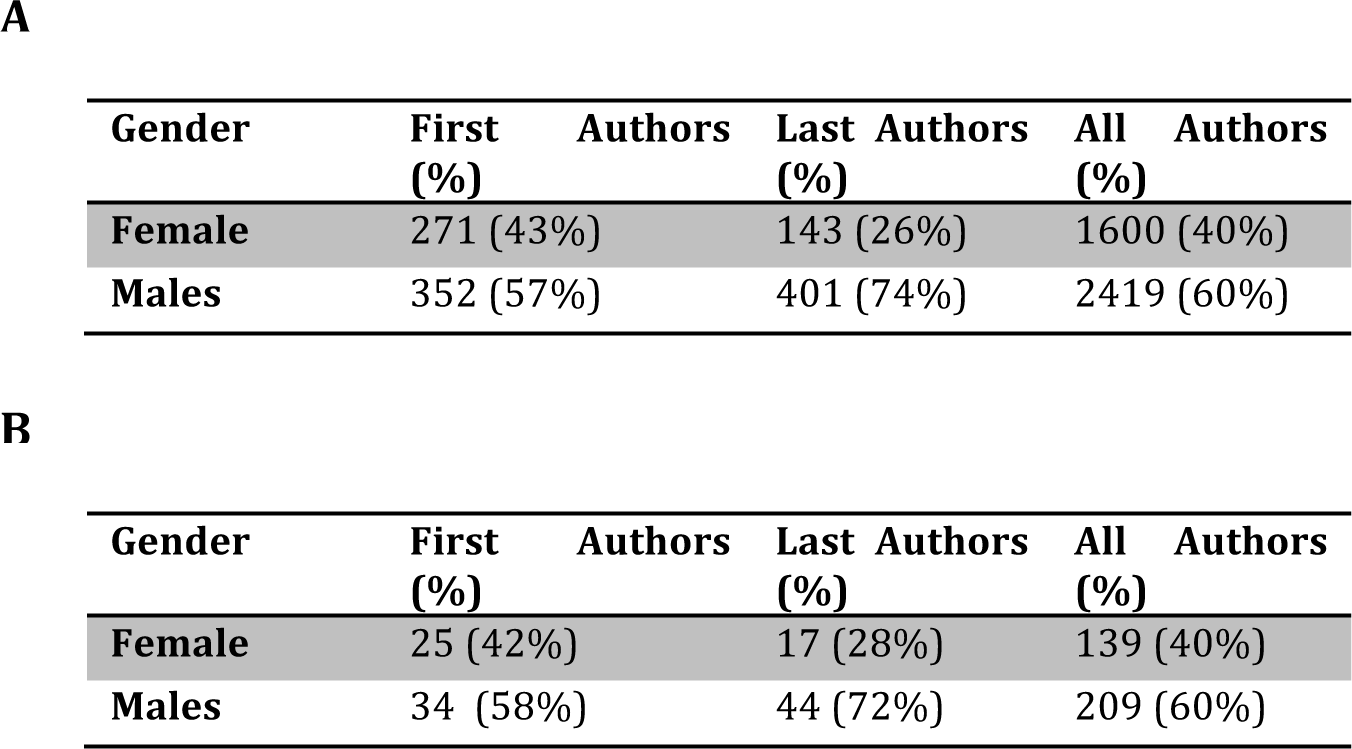
Gender composition of authors in genome editing (2017) based on about 50% of first names where gender could be assigned. **(A)** CRISPR and **(B)** TALENs based genome editing.

**Table 2:**
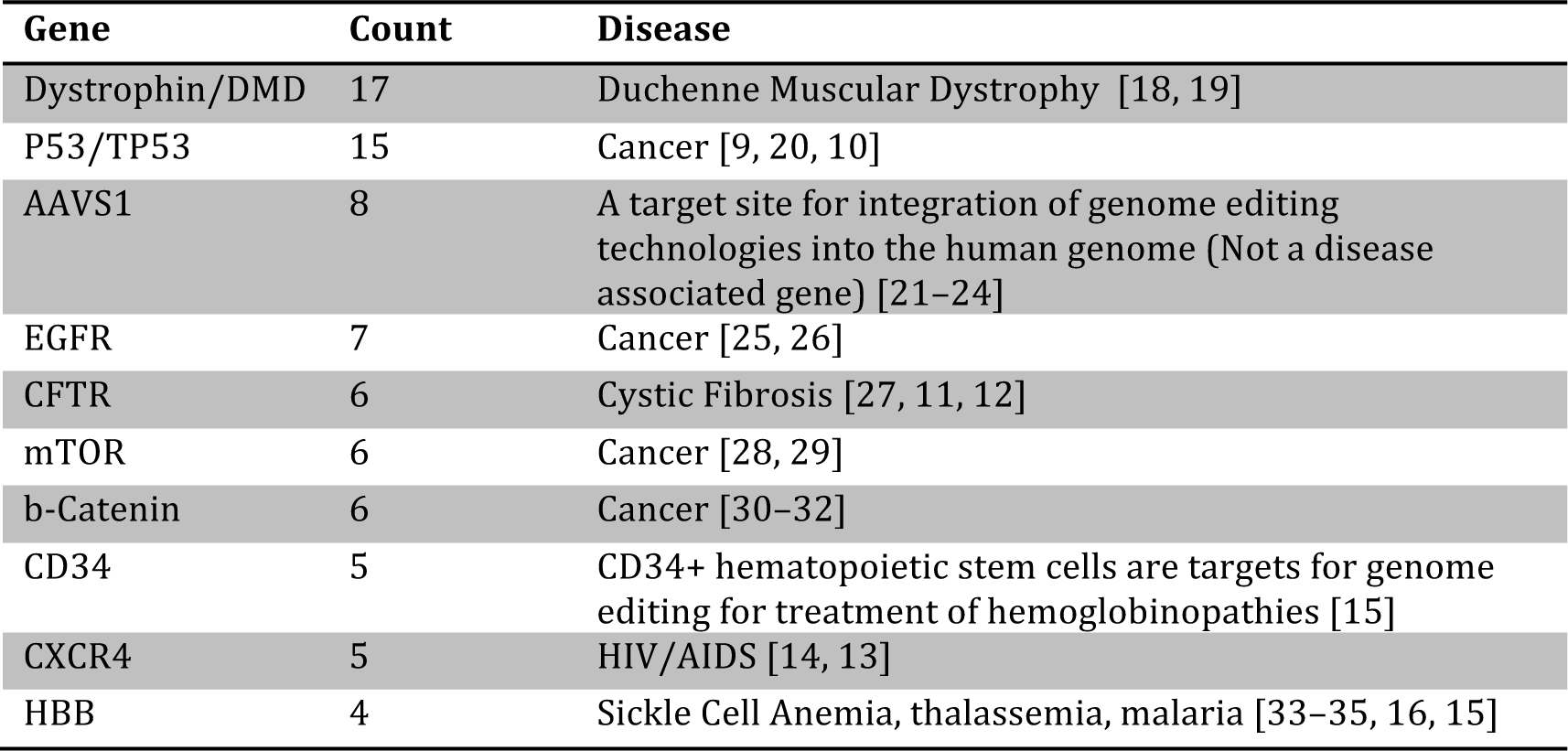
Top genes and diseases associated with CRISPR genome editing studies in 2017.

| Gene | Count | Disease |
| --- | --- | --- |
| Dystrophin/DMD | 17 | Duchenne Muscular Dystrophy [18, 19] |
| P53/TP53 | 15 | Cancer [9, 20, 10] |
| AAVS1 | 8 | A target site for integration of genome editing technologies into the human genome (Not a disease associated gene) [21–24] |
| EGFR | 7 | Cancer [25, 26] |
| CFTR | 6 | Cystic Fibrosis [27, 11, 12] |
| mTOR | 6 | Cancer [28, 29] |
| b-Catenin | 6 | Cancer [30–32] |
| CD34 | 5 | CD34+ hematopoietic stem cells are targets for genome editing for treatment of hemoglobinopathies [15] |
| CXCR4 | 5 | HIV/AIDS [14, 13] |
| HBB | 4 | Sickle Cell Anemia, thalassemia, malaria [33–35, 16, 15] |

Another alternative approach for identifying diseases being targeted by genome editing technologies is using named entity recognition to extract disease names directly from abstracts. When this approach was applied to screen abstracts, the top diseases for CRISPR genome editing were cancer/tumor/carcinoma (355 articles), Duchenne Muscular Dystrophy (DMD, 24), malaria (14), HIV/AIDS (18) and hepatitis B/C virus (13). On the other hand, top diseases for TALENs were cardiovascular diseases (9 articles), HIV/AIDS (9), hepatitis B/C virus (8) and hemoglobinopathies (4 articles).

Interestingly, malaria, a disease that predominantly affects sub-saharan Africa is one of the top diseases for CRISPR research although researchers from the region are heavily underrepresented in the field. While Africa is underrepresented in several areas of research besides genome editing, underrepresentation in genome editing is crucial as the technology has a great potential in malaria eradication but also poses broader ethical issues [36, 37].

An understanding of species that are research targets of genome editing technologies could help identify neglected species by the emerging technology. Therefore, we extracted words referring to various species from the abstracts. Among the top species of interest in genome editing research using CRISPR, humans were the most common (519 articles), followed by mice (306), yeast (*S. cerevisiae*, 76), zebrafish (60), rice (40), pigs (34), drosophila (32), HIV (28), and *Caenorhabditis elegans* (27). Similarly, for TALENs, human cells were the most commonly studied (38 articles), mice (22), zebrafish (10), rat (10), pigs (9), rice (5) and wheat (5).

## Discussion

Genome editing technologies could revolutionize treatment of several diseases and open new ways for engineering biological organisms for medical and non-medical applications. An understanding of the global distribution of genome editing research could inform policy development, funding strategies and identify research biases that could lead to disparities in the application of the technology. Unfortunately, with thousands of papers being published in the field, manual curation of their content is not optimal. Automated approaches to mining literature could fill this gap by providing an unbiased and rapid assessment of progress in the field. However, there are several drawbacks to applying automated literature-mining approaches. First, mining of published literature cannot capture all research. For example, proprietary research in commercial companies is often not published. Furthermore, even in academia, research may not always be published immediately. Secondly, many publications are behind paywalls thereby limiting literature mining to only abstracts. Finally, computational approaches for mining text using natural language processing (NLP) and machine learning are still not able to cope with the complexity ambiguity and diversity of human language.

In this work, literature mining of genome editing publications shows that the development of the technology disproportionately occurs in the US and China. While the US leads in CRISPR based studies, China leads in TALENs. Thus, these two nations may potentially be the first to see significant benefits of genome editing technologies. While today no clinical applications have been approved for genome editing technologies, the technology could provide early economic benefits such as new careers especially for the researchers needed to develop the technology. It is conceivable that regions of the world already leading research in this area will be well positioned with the human capacity required to advance it which will result in disproportionate economic gains. Regions of the world such as Africa that are underrepresented in research in this field will left behind and may not be able to catch up.

Genome editing research is not immune to entrenched biases that exist in our society today. In particular, gender biases in science and technology could bias the development of genome editing technologies. However, the relatively recent emergence of the field compared to other areas of technology provides a unique window of opportunity that should be used to avoid new biases and address those inherited from related fields. The results presented in this work show that while there are 60% male and 40% female researchers in this field based on publications, the principal investigators are predominantly male (76%) which is consistent with observations that women are even more underrepresented in leadership roles in Science, Technology, Engineering, Mathematics and Medicine (STEMM) fields [38].

The disproportionate development of genome editing technologies towards a select number of diseases, genes and animals or plants is also notable. It is encouraging that among the top diseases where genome editing research is ongoing are rare diseases such as Duchenne Muscular Dystrophy [19] and those that predominantly afflict some of the poorest regions of the world such as malaria [37, 39]. Unfortunately, in the case of malaria, researchers based in areas affected by the disease (sub-Saharan Africa) are highly underrepresented in genome editing research. This misalignment needs to be addressed because objective assessment of the safety, ethical and economic issues arising from genome editing technologies will require regulatory approvals by local experts and will directly impact local communities. Furthermore, a significant number of genome editing publications are in subscription journals and may be inaccessible to researchers in poorer regions of the world making it harder for them to compete in the field. Proactive engagement of researchers in areas where field applications such as mosquito gene drives will be deployed occur using genome editing technologies will be crucial [36]. One such partnership is the Target Malaria consortium that brings together various stakeholders including researchers and risk-assessment specialists from Europe and Africa [40].

Genome editing is a powerful technology. Its disproportionate development by a few researchers, regions of the world or in a gender-biased manner will limit its full potential. Furthermore, its disproportionate application to select genes, diseases and organisms could limit its capacity as an aid for deepening our understanding of basic biological processes across various organisms, engineering new biological systems for food or industrial purposes and eventually finding cures to several diseases.

## Materials and Methods

### Data retrieval

To download abstracts of publications on genome editing, the PubMed literature database was queried on May 21^st^ 2018 using the *‘easyPubMed’* package in R. Briefly, the PubMed database was queried using 4 search queries independently, two for CRISPR/Cas based genome editing and two for TALENs based genome editing. To obtain genome editing publications involving CRISPR/Cas systems in 2017 the in the search term used was “CRISPR genome editing AND (2017[PDAT])” while to download genome editing publications involving CRISPR/Cas systems in 2018, the search term used was “CRISPR genome editing AND (2018[PDAT])”. Abstracts on TALENs based genome editing were downloaded using the search terms “TALENs genome editing AND (2017[PDAT])” for publications in 2017 and “TALENs genome editing AND (2018[PDAT])” for publications in 2018. Because CRISPR research does not always involve genome editing, these numbers capture only the subset of papers where the abstracts explicitly mention genome editing. Abstracts were downloaded in XML format to allow further retrieval of metadata such as author names, institutional affiliations, addresses and countries.

### Analysis of author names, gender and institutional affiliations

Author names, journals and institutional addresses were directly extracted from the downloaded XML abstracts using the R package *‘easyPubMed’*. Authors were assigned a gender using data from the US Social Security Administration baby names as implemented in the R package *‘gender’*. Institutional addresses were further processed to identify country names using the R package *‘map’*.

### Named entity recognition to identify diseases, gene names and species

Extraction of named biomedical entities such as genes, diseases and species referred to in the downloaded abstracts was performed using the R package *‘pubmed.mine.R’* [41].

## Supplementary Materials

**Supplementary Table 1:** Number of articles per journal in the top 50 journals for CRISPR genome editing in 2017.

| Journal Name | Number of articles |
| --- | --- |
| Sci Rep | 81 |
| Methods Mol. Biol. | 49 |
| Nat Commun | 32 |
| Nature | 30 |
| PLoS ONE | 27 |
| Proc. Natl. Acad. Sci. U.S.A. | 22 |
| Front Plant Sci | 21 |
| ACS Synth Biol | 18 |
| Mol. Cell | 18 |
| Nat. Biotechnol. | 18 |
| Nucleic Acids Res. | 18 |
| Plant Biotechnol. J. | 18 |
| Prog Mol Biol Transl Sci | 18 |
| Mol Ther Nucleic Acids | 17 |
| Oncotarget | 17 |
| Methods | 15 |
| Sheng Wu Gong Cheng Xue Bao | 15 |
| Mol. Ther. | 14 |
| Adv. Exp. Med. Biol. | 13 |
| Science | 13 |
| Sci China Life Sci | 12 |
| Mamm. Genome | 11 |
| ACS Chem. Biol. | 10 |
| Int J Mol Sci | 10 |
| Yale J Biol Med | 10 |
| Cell | 9 |
| Circ. Res. | 9 |
| G3 (Bethesda) | 9 |
| J Genet Genomics | 9 |
| J. Biotechnol. | 9 |
| J. Cell. Biochem. | 9 |
| Stem Cell Reports | 9 |
| Cell Rep | 8 |
| Crit. Rev. Biotechnol. | 8 |
| Dev. Biol. | 8 |
| Elife | 8 |
| Genome Biol. | 8 |
| J Vis Exp | 8 |
| mSphere | 8 |
| Plant Cell Rep. | 8 |
| Stem Cell Res | 8 |
| Transgenic Res. | 8 |
| Bio Protoc | 7 |
| Biotechnol. Bioeng. | 7 |
| Brief Funct Genomics | 7 |
| Cell Stem Cell | 7 |
| Curr Issues Mol Biol | 7 |
| Microb. Cell Fact. | 7 |
| Mol Plant | 7 |
| Mol. Biol. Cell | 7 |
| PLoS Genet. | 7 |
| Protein Cell | 7 |

**Supplementary Table 2:** Number of articles per journal in the top journals for TALENs genome editing in 2017.

| Journal Name | Number of Papers |
| --- | --- |
| Methods Mol. Biol. | 12 |
| Front Plant Sci | 6 |
| PLoS ONE | 5 |
| Dev. Biol. | 4 |
| Mol Ther Methods Clin Dev | 4 |
| Prog Mol Biol Transl Sci | 4 |
| Crit. Rev. Biotechnol. | 3 |
| Hum. Gene Ther. | 3 |
| ACS Synth Biol | 2 |
| Cell Biosci | 2 |
| Cell. Mol. Life Sci. | 2 |
| Front Mol Neurosci | 2 |
| J. Biotechnol. | 2 |
| Nat Protoc | 2 |
| Nucleic Acids Res. | 2 |
| Sci China Life Sci | 2 |
| Sci Rep | 2 |
| Sheng Wu Gong Cheng Xue Bao | 2 |

**Supplementary Table 3:**
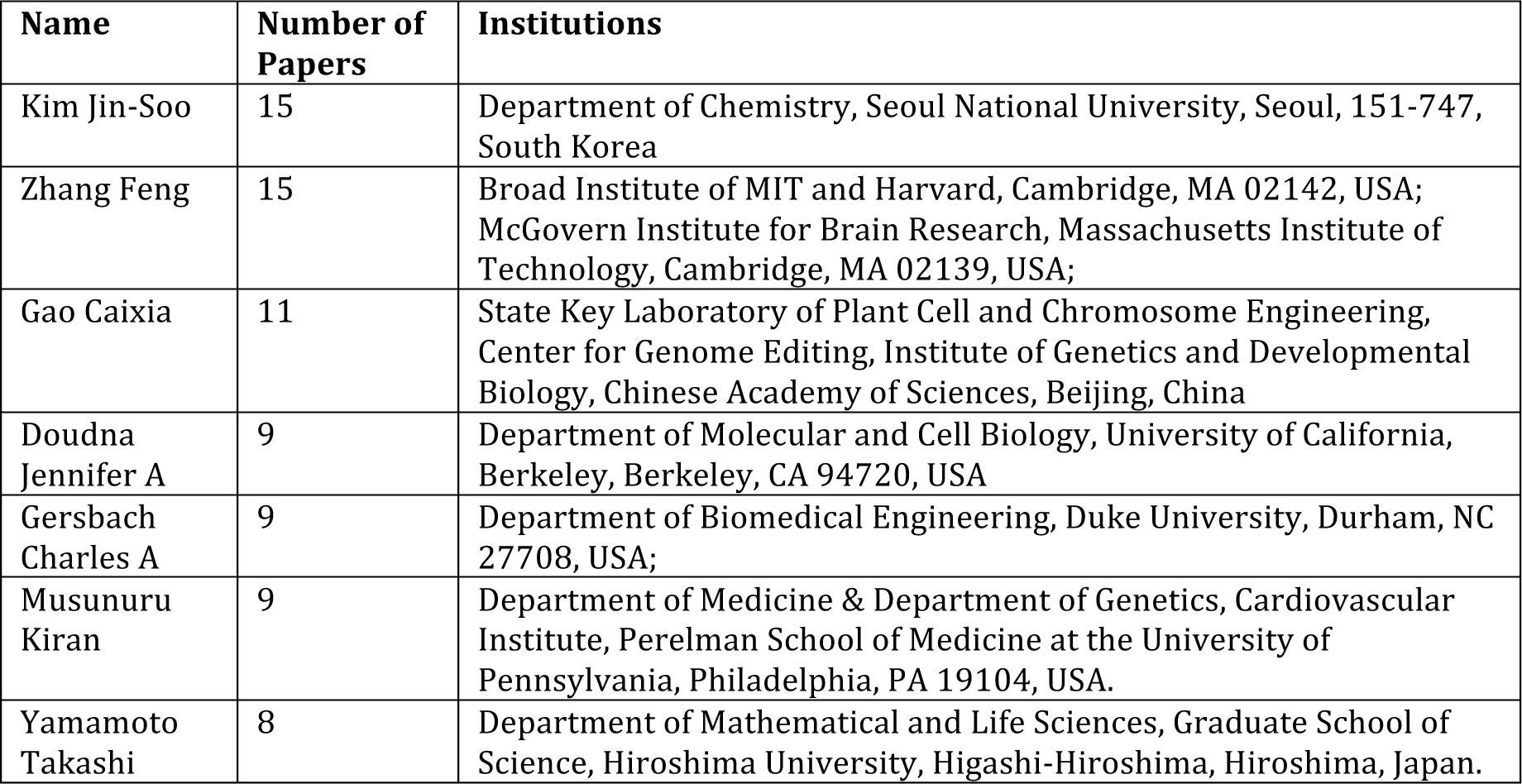
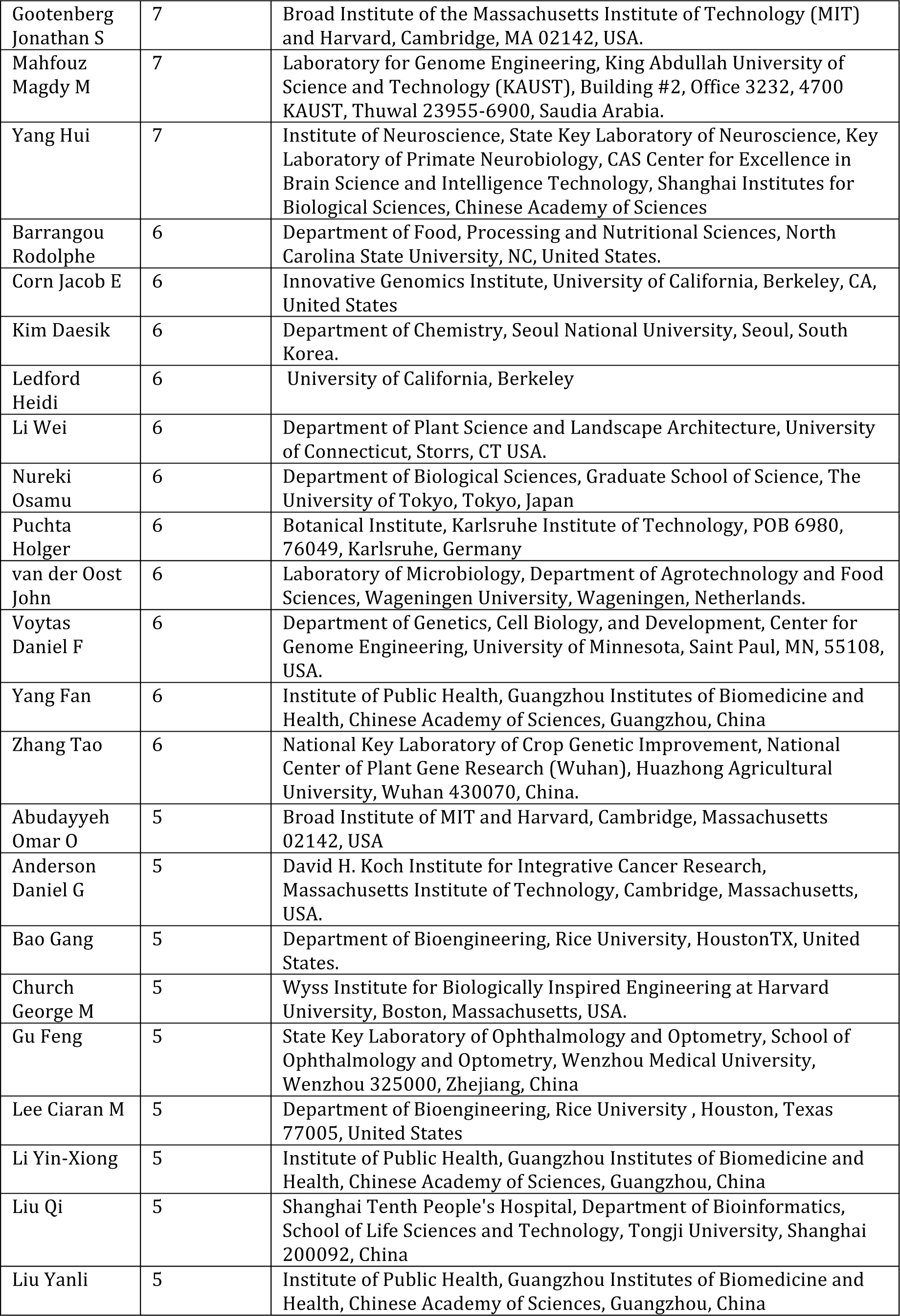
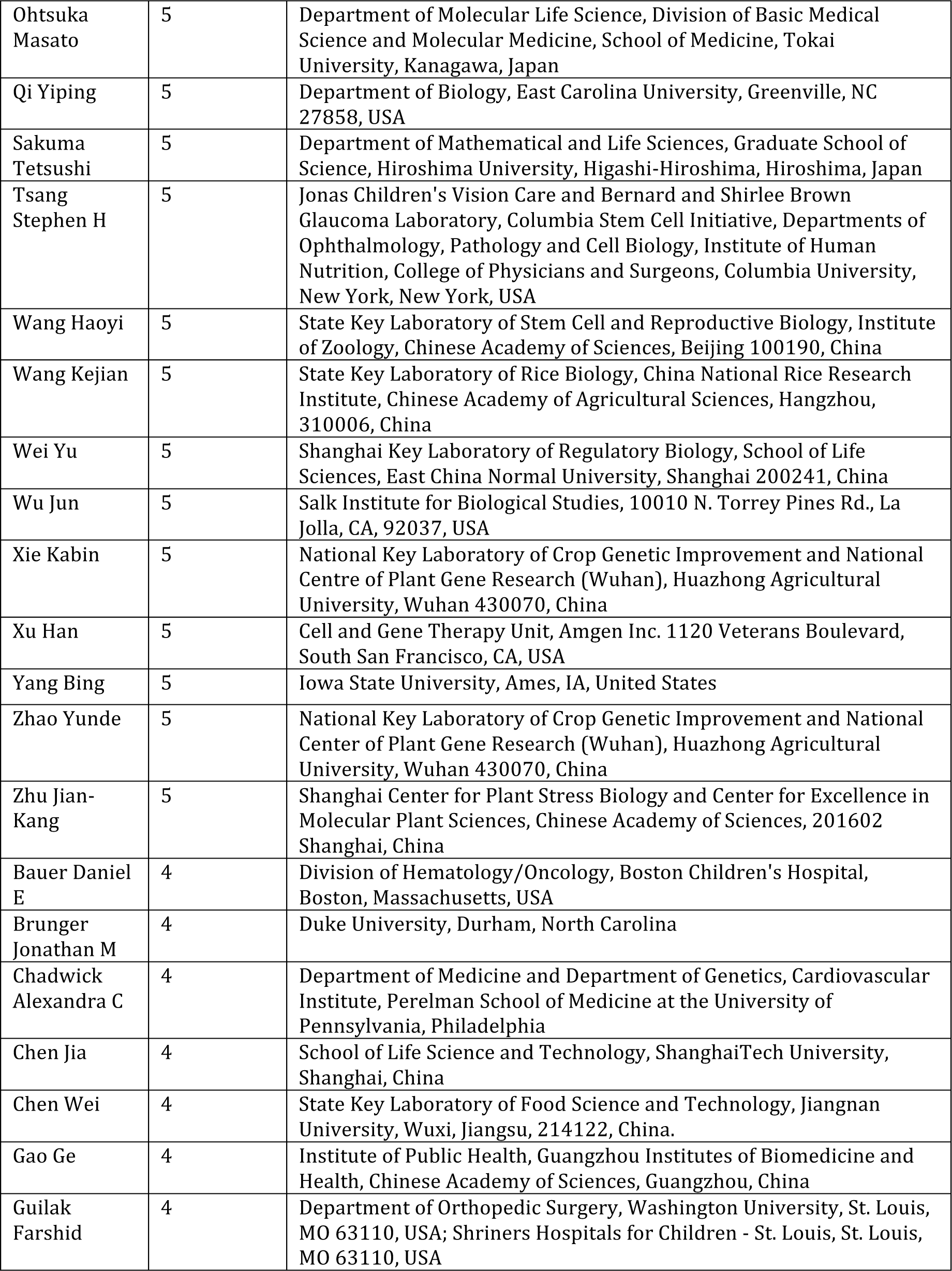
Author names and publication count for the top 50 authors for CRISPR (2017).

**Supplementary Table 4:** Author names and publication count for the top 50 authors for TALENs (2017).

| **Names** | **Number of Papers** | **Institution** |
| --- | --- | --- |
| Sakuma Tetsushi | 5 | Department of Mathematical and Life Sciences, Graduate School of Science, Hiroshima University, Hiroshima, Japan |
| Yamamoto Takashi | 5 | Department of Mathematical and Life Sciences, Graduate School of Science, Hiroshima University, Hiroshima, Japan |
| Cathomen Toni | 4 | Institute for Transfusion Medicine and Gene Therapy, Medical Center- University of Freiburg, Freiburg, Germany; Faculty of Medicine, University of Freiburg, Freiburg, Germany. |
| Mussolino Claudio | 4 | Institute for Cell and Gene Therapy, University Medical Center Freiburg, Freiburg, Germany |
| Alzubi Jamal | 2 | Institute for Transfusion Medicine and Gene Therapy, Medical Center - University of Freiburg, Freiburg, Germany |
| Bortesi Luisa | 2 | Institute for Molecular Biotechnology, RWTH Aachen University, Worringerweg 1, 52074 Aachen, Germany |
| Fischer Rainer | 2 | Institute for Molecular Biotechnology, RWTH Aachen University, Worringerweg 1, 52074 Aachen, Germany |
| Germini Diego | 2 | UMR 8126, Université Paris Sud - Paris Saclay, CNRS, Institut Gustave Roussy, 94805 Villejuif, France |
| Gu Feng | 2 | State Key Laboratory of Ophthalmology and Optometry, School of Ophthalmology and Optometry, Wenzhou Medical University, Wenzhou 325000, Zhejiang, China |
| He Xiubin | 2 | State Key Laboratory of Ophthalmology, Optometry and Vision Science, School of Ophthalmology and Optometry, Eye Hospital, Wenzhou Medical University, Wenzhou, China |
| Hendel Ayal | 2 | The Mina and Everard Goodman Faculty of Life Sciences and Advanced Materials and Nanotechnology Institute, Bar-Ilan University, Ramat-Gan 52900 Israel |
| Horb Marko E | 2 | National Xenopus Resource and Eugene Bell Center for Regenerative Biology and Tissue Engineering, Marine Biological Laboratory, Woods Hole, MA 02543, United States |
| Kaneko Takehito | 2 | Institute of Laboratory Animals, Graduate School of Medicine, Kyoto University, Kyoto 606-8501, Japan |
| Karakikes Ioannis | 2 | Stanford Cardiovascular Institute, Stanford University School of Medicine, Stanford, CA, 94305, USA. |
| Koller Ulrich | 2 | EB House Austria, Research Program for Molecular Therapy of Genodermatoses, Department of Dermatology, University Hospital of the Paracelsus Medical University, Salzburg, Austria |
| Musunuru Kiran | 2 | Cardiovascular Institute, Department of Medicine, Perelman School of Medicine at the University of Pennsylvania, Philadelphia, PA, USA |
| Reichelt Julia | 2 | EB House Austria, Research Program for Molecular Therapy of Genodermatoses and Department of Dermatology, University Hospital of the Paracelsus Medical University Salzburg, 5020 Salzburg, Austria |
| Sasakura Yasunori | 2 | Shimoda Marine Research Center, University of Tsukuba, Shimoda, Shizuoka 415-0025, Japan |
| Seeger Timon | 2 | Stanford Cardiovascular Institute, Stanford University School of Medicine, Stanford, CA, 94305, USA. |
| Sjakste Nikolajs | 2 | Faculty of Medicine, University of Latvia, Riga, Latvia. |
| Termglinchann Vittavat | 2 | Stanford Cardiovascular Institute, Stanford University School of Medicine, Stanford, CA, 94305, USA |
| Thrasher Adrian J | 2 | Institute of Child Health, University College London, London, United Kingdom |
| Treen Nicholas | 2 | Lewis-Sigler Institute for Integrative Genomics, Princeton University, Princeton, NJ, USA |
| Tsfasman Tatiana | 2 | UMR 8126, Université Paris Sud - Paris Saclay, CNRS, Institut Gustave Roussy, 94805 Villejuif, France |
| Vassetzky Yegor | 2 | UMR8126, Université Paris Sud - Paris Saclay, CNRS, Institut Gustave Roussy, 94805 Villejuif, France |
| Wang Haoyi | 2 | State Key Laboratory of Stem Cell and Reproductive Biology, Institute of Zoology, Chinese Academy of Sciences, Beijing 100101, China |
| Wu Joseph C | 2 | Department of Medicine, Division of Cardiovascular Medicine, Stanford University School of Medicine, Stanford, California 94305, USA |
| Yoshida Keita | 2 | Shimoda Marine Research Center, University of Tsukuba, Shimoda, Shizuoka, Japan |

